# DNA barcoding British *Euphrasia* reveals deeply divergent polyploids but lack of species-level resolution

**DOI:** 10.1101/164681

**Authors:** Xumei Wang, Galina Gussarova, Markus Ruhsam, Natasha de Vere, Chris Metherell, Peter M. Hollingsworth, Alex D. Twyford

## Abstract

**Background and aims:** DNA barcoding is emerging as a useful tool not only for species identification but for studying evolutionary and ecological processes. Although plant DNA barcodes do not always provide species-level resolution, the generation of large DNA barcode datasets can provide insights into the mechanisms underlying the generation of species diversity. Here, we use DNA barcoding to study evolutionary processes in taxonomically complex British *Euphrasia*, a group with multiple ploidy levels, frequent self- fertilization, young species divergence and widespread hybridisation.

**Methods:** We sequenced the core plant barcoding loci, supplemented with additional nuclear and plastid loci, in representatives of all 19 British *Euphrasia* species. We analyse these data in a population genetic and phylogenetic framework. We then date the divergence of haplotypes in a global *Euphrasia* dataset using a time-calibrated Bayesian approach implemented in BEAST.

**Key results:** No *Euphrasia* species has a consistent diagnostic haplotype. Instead, haplotypes are either widespread across species, or are population specific. Nuclear genetic variation is strongly partitioned by ploidy levels, with diploid and tetraploid British *Euphrasia* possessing deeply divergent ITS haplotypes (D_XY_ = 5.1%), with haplotype divergence corresponding to the late Miocene. In contrast, plastid data show no clear division by ploidy, and instead reveal weakly supported geographic patterns.

**Conclusions:** Using standard DNA barcoding loci for species identification in *Euphrasia* will be unsuccessful. However, these loci provide key insights into the maintenance of genetic variation, with divergence of diploids and tetraploids suggesting that ploidy differences act as a barrier to gene exchange in British *Euphrasia*, with rampant hybridisation within ploidy levels. The scarcity of shared diploid-tetraploid ITS haplotypes supports the polyploids being allotetraploid in origin. Overall, these results show that even when lacking species-level resolution, DNA barcoding can reveal insightful evolutionary patterns in taxonomically complex genera.

## INTRODUCTION

DNA barcoding is a valuable tool for discriminating among species, and these data often give insights into identity that are overlooked based on morphology alone (Hebert and Gregory, 2005). Plant DNA barcoding has been used for species discovery, reconstructing historical vegetation types from frozen sediments, surveying environmental variation, and many other research topics (reviewed in Hollingsworth et al., 2016). Despite the extensive uptake of DNA barcoding, there are numerous reports of taxon groups where DNA barcodes do not provide exact plant species identification, and where DNA barcode sequences are shared among related species (Percy et al., 2014, Yan et al., 2015a, Yan et al., 2015b, Spooner, 2009). Identifying the causes of why DNA barcoding ‘fails’ is essential to help guide the development of future DNA barcoding systems. If species discrimination is usually limited by information content of the core barcoding loci, then improvements can be made by extending the DNA barcode to harness variation across the whole plastid genome. In contrast, if limits to species discrimination are caused by hybridisation, incomplete lineage sorting, and polyploidy, then we may see no improvement with more plastid data, and future barcoding systems will need to target the nuclear genome, as well as use new analytical methods to cope with many independent loci (Coissac et al., 2016, Hollingsworth et al., 2016). As such, more genetic studies of complex taxa groups are required in order to identify the underlying evolutionary processes that can cause DNA barcoding to fail. In addition, the generation of large data sets of DNA sequence data from multiple individuals of multiple species in such groups also provide datasets that can shed light onto evolutionary relationships and evolutionary divergence, without a need for the barcode markers to track species boundaries.Postglacial species radiations of taxonomically complex groups in Northern Europe are a case where we may not expect a clear cut-off between intraspecific variation and interspecific divergence and thus DNA barcoding may provide limited discriminatory power. Despite this, DNA barcoding may still be valuable if used to identify evolutionary and ecological processes that result in shared sequence variation. For example, many postglacial taxa are characterised by a combination of: (1) recent postglacial speciation, (2) extensive hybridisation, (3) frequent self-fertilization, (4) divergence involving polyploidy. Our expectation is that factors 1 + 2 will cause DNA barcode sequences to be shared among geographically proximate taxa, while 3 will cause barcodes to be population rather than species-specific (Hollingsworth et al., 2011). Factor 4, polyploidy, will manifest as shared haplotypes between recent polyploids and their parental progenitors, or deep haplotype divergence in older polyploid groups, where ploidy act as a reproductive isolating barrier and allows congeneric taxa to accumulate genetic differences. Many of these interacting factors are common across taxonomically complex postglacial groups (Ennos, 2005), which include the *Arabidopsis arenosa* complex (Schmickl et al., 2012), *Cerastium* (Brysting et al., 2007), *Epipactis* (Squirrell et al., 2002) and *Galium* (Kolář et al., 2015). As such, the application of DNA barcoding to such groups will not only reveal the prevalence of haplotype sharing and the potential for species discrimination, but improve our understanding of processes underlying shared variation.

One example of a taxonomically challenging group showing postglacial divergence is British *Euphrasia* species. This group of 19 taxa are renowned for their difficult species identification, and at present only a handful of experts can identify these species in the field. Morphological species identification is difficult due to their small stature, combined with species being defined by a complex suite of overlapping characters (Yeo, 1978). They are also generalist hemiparasites and thus phenotypes are plastic and depend upon host quality (Svensson and Carlsson, 2004). DNA barcode-based identification could partly resolve these identification issues, and lead to a greater understanding of species diversity and distributions in this under-recorded group. This is particularly important as a number of *Euphrasia* species are critically rare and of conservation concern, while others are ecological specialists that are useful indicators of habitat type (French et al., 2008). More generally, DNA barcoding data could reveal the processes structuring genetic diversity and those that are responsible for recent speciation.

Previous broad-scale surveys of *Euphrasia* using AFLP and microsatellites have shown a significant proportion of genetic variation is partitioned by ploidy groups and by species, despite extensive hybridisation (French et al., 2008). Here, we follow-on from previous population genetic studies by testing the utility of DNA barcoding across British *Euphrasia*. Our first aim is to assess whether DNA barcoding is informative for species recognition in this young postglacial group. Our second aim is to understand the evolutionary factors that may explain patterns of shared DNA sequences. We first sequence British populations for the standard DNA barcoding loci, and analyse the distribution of haplotypes across populations and species. To better understand the historical context of haplotype sharing we sequence additional loci and use dated phylogenetic analyses to infer divergence ages. Overall, these results are used to understand the efficacy of genetic tools for studying species-level variation in a taxonomically complex group.

## MATERIALS AND METHODS

### Specimen sampling

The 19 currently recognised British *Euphrasia* species are all annuals, selfers or mixed-mating small herbaceous plants, which occur in a range of habitats including coastal turf, chalk downland, mountain ridges and heather moorland. The species can be divided into two groups, glabrous or short-eglandular hairy tetraploids (15 species, Fig 1A), or long glandular hairy diploids (4 species, Fig. 1B). Our sampling included representatives of all British species, with samples collected from across widespread populations (Fig. 1c, Table S1, [**Supplementary Information**]). Samples were collected in South West England and Wales to allow us to include mixed populations of diploids and tetraploids, early generation diploid x tetraploid hybrids, and two diploid hybrid species hypothesised to be derived from diploid x tetraploid crosses (*E. vigursii*, parentage: *E. rostkoviana* x *E. micrantha*), *E. rivularis* (parentage: *E. anglica* x *E. micrantha*; Yeo, 1956). Samples from Scotland allowed us to sample complex tetraploid taxa and tetraploid hybrids, plus scarcer Scottish diploids. Our sampling scheme investigated range-wide variation by targeting many taxa and populations, but without intrapopulation sampling. This is because prior work has shown low intrapopulation diversity, with populations frequently fixed for single haplotypes (French et al., 2008). All samples were identified by *Euphrasia* experts Alan Silverside or Chris Metherell.

**Figure 1.**
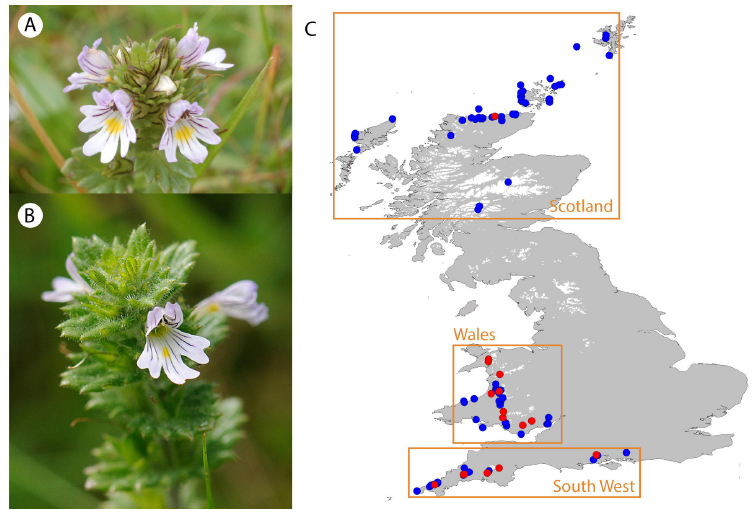
*Euphrasia* samples used in this study. (A) Tetraploid British *Euphrasia* (here *E. arctica*) have glabrous leaves sometimes with sparse short eglandular hairs or bristles. (B) Diploid British *Euphrasia* have long glandular hairs. (C) Collection sites of *Euphrasia* DNA samples. Diploids are shown in red, tetraploids in blue. Orange boxes correspond to the three broad sampling areas.

For our population-level DNA barcoding study, we analysed a total of 133 individuals, with 106 samples representing 19 species, as well as 27 samples from 14 putative hybrids. We sequenced samples for the core plant DNA barcoding loci, *mat*K and *rbc*L (Hollingsworth et al., 2009), as well as ITS2 (China Plant BOL Group, 2011), and supplemented these loci with *rpl32-trnL*^*UAG*^, which has been informative in prior population studies of *Euphrasia* (Stone, 2013).

For our broader molecular phylogenetic and molecular dating analysis, we expanded the sampling in the molecular phylogeny of the genus by Gussarova et al. (2008), to include a detailed sample of British taxa. The previous analysis included 40 taxa for the nuclear ribosomal internal transcribed spacer (ITS), and 50 taxa for plastid DNA (Gussarova et al., 2008). We sequenced samples to match the previous data matrix, which included: the *trnL* intron (Taberlet et al., 1991), intergenic spacers *atpB-rbcL* (Hodges and Arnold, 1994) and *trnL-trnF* (Taberlet et al., 1991), and *ITS* (White et al. 1990).

### DNA extraction, PCR amplification, and sequencing

DNA was extracted from silica dried tissue using the DNeasy Plant Mini kit (Qiagen, Hilden, Germany) following the manufacturer's protocol, but with an extended incubation of 1 hour at 65 °C. These DNA samples were added to existing DNA extractions from 68 individuals from French et al. (2008).

PCRs were done in two laboratories following separate protocols. We applied the following conditions for most taxa. We performed PCRs in 10 μL reactions, with DNA amplification and PCR conditions for each primer given in Table S2 [**Supplementary Information**]. We visualised PCR products on a 1% agarose gel, with 5 μL of PCR product purified for sequencing with ExoSAP-IT (USB Corporation, Cleveland, OH, USA) using standard protocols. Sequencing was performed in 10 μL reactions containing 1.5 μL 5 × BigDye buffer (Life Technologies, Carlsbad, CA, USA), 0.88 μL BigDye enhancing buffer BD × 64 (MCLAB, San Francisco, CA, USA), 0.125 μL BigDye v3.1 (Life Technologies), 0.32 μM primer and 1 μM of purified PCR product. We sequenced PCR products on the ABI 3730 DNA Analyser (Applied Biosystems, Foster City, CA, USA) at Edinburgh Genomics. A subset of sequences were generated as part of the effort to DNA barcode the UK Flora, and followed a different set of protocols, detailed in de Vere et al. (2012). We assembled, edited and aligned sequences using Geneious v. 8 (Biomatters, Auckland, New Zealand), with manual editing. Indels were coded as unordered binary characters and appended to the matrices. We used gap coding as implemented in Gapcoder (Young and Healy, 2003), with indels treated as point mutations and equally weighted with other mutations.

### Population genetic analyses of British samples

We examined patterns of sequence variation using a range of population genetic methods. We investigated the amount of sequence diversity in these recently diverging species using descriptive statistics, and then tested the cohesiveness of taxa using analysis of molecular variance (AMOVA) and related methods. Analyses were performed separately on ITS2 and a concatenated matrix of plastid data. For plastid data, haplotypes were determined from nucleotide substitutions and indels of the aligned sequences. Basic population genetic statistics were performed in Arlequin Version 3.0 (Excoffier and Lischer, 2010), and this included the number of haplotypes, as well as hierarchical AMOVA in groups according to:(1) ploidy levels (diploid vs tetraploid); (2) geographic regions (Wales, England, Scotland);(3) species. AMOVAs were performed on all taxa, and repeated for ploidy levels and geographic regions on a dataset only including confirmed species (i.e. excluding hybrids). Sequence diversity and divergence statistics were estimated with DnaSP (Librado and Rozas, 2009), which included: average nucleotide diversity across taxa (Pi), Watterson’s theta (per site), Tajima’s D and divergence between ploidy levels (D_XY_).

Genetic divergence among sampling localities were explored with Spatial Analysis of Molecular Variance (SAMOVA; Dupanloup et al., 2002), implemented in SPADS v.1.0 (Dellicour and Mardulyn, 2014). SAMOVA maximises the proportion of genetic variance due to differences among populations (F_CT_) for a given number of genetic clusters (*K*-value). We considered the best grouping to have the highest F_CT_ value after 100 repetitions. This analysis investigated interspecific differentiation, thus only used species samples, excluding hybrids.

The relationships between haplotypes was inferred by constructing median-joining networks (MJM; Bandelt et al., 1999) with the program NETWORK v.4.6.1.1 (available at http://www.fluxus-engineering.com/), treating gaps as single evolutionary events.

### Phylogenetic analyses and molecular dating

We used Bayesian phylogenetic analyses in MrBayes v. 3.1.2 (Huelesenbeck & Ronquist 2001) to infer species relationships and broad-scale patterns of colonisation. Our analyses used a sequence matrix that included our newly sampled British taxa in addition to previous global *Euphrasia* samples from Gussarova et al. (2008). We selected the best fitting model of nucleotide substitution using the Akaike Information Criterion (AIC) with an empirical correction for small sample sizes implemented in MrAIC (Nylander, 2004). Using GTR +G as the best model for the plastid dataset and SYM + G for the ITS dataset we ran two sets of four Markov Chain Monte Carlo (MCMC) runs for 5,000,000 generations. Indels were included as a separate partition with a restriction site (binary) model. We sampled every 1000^th^ generation and used a burnin of 2,500,000, and default priors. We confirmed chain convergence by observing the average standard deviation of split frequencies and by plotting parameter values in Tracer v. 1.6 (Rambaut & Drummond 2013).

We used BEAST v. 1.8.2 (Drummond et al. 2012) to estimate the divergence ages of major *Euphrasia* lineages occurring in Britain. This analysis used our British samples in conjunction with the full global dataset of *Euphrasia* (Gussarova et al., 2008). Our analysis gives crude dates due to the lack of available fossils for calibration, but these estimates allow us to compare between very recent (postglacial) divergence, and much older divergence events. We only analysed ITS sequences, due to the lack of support obtained for the plastid phylogeny (see results). The analysis included two partitions: one containing all ITS sequences (638 bp) and the other containing 38 gap-coded indels. We applied the same substitution models as those used in the MrBayes analyses. For the binary characters, we used a stochastic Dollo model. Separate analyses were run testing for the strict vs the uncorrelated lognormal (UCLN) molecular clock. The branching was modelled using the Yule tree prior, which assumes a constant speciation rate per lineage. The root age was set with a normal prior of mean = 2.65×10^7^ and SD = 5×10^5^, according to the results obtained based on a calibrated phylogeny of *Euphrasia* by Gussarova et al. (2008). This prior analysis used geological dating of volcanic islands to set priors as maximum ages for colonization of endemic *Euphrasia* species. Substitution rate priors were based on Key et al. (2006), ranging from 0.38 ×10^-^9^^ to 8.34 ×10^-^9^^ substitutions/site/year. Data analyses were run for 50,000,000 generations, logging every 50,000^th^ generation. We compared the fit of the clock models using AICM criterion implemented in Tracer. The AICM values obtained were very similar between the two models: 8723.681+/-0.685 vs. 8882.911+/-1.582. The strict clock was chosen as it is a simpler model and with lower AICM. These phylogenetic results were compared with those from an empty alignment using the same values for priors.

## RESULTS

### ITS haplotype diversity in British populations

The final ITS2 alignment contained 130 individuals representative of all British taxa, and was 380 bp in length. Only two samples (of *E. scottica*) presented double peaks, and were excluded from analysis. Overall diversity across taxa was modest, with a nucleotide diversity (Pi) of 2.3%, and theta (per site) of 0.01781. There were 33 nucleotide substitutions and one indel, from which we called 23 haplotypes. All sequences have been deposited in GenBank (Accession numbers given on acceptance).

The ITS haplotypes revealed strong partitioning by ploidy. Of the 23 haplotypes, three (H1, H20 and H21) were restricted to diploids, and 19 to tetraploids, with only one haplotype (H2) shared across ploidy levels (Table 1). Haplotype H2 was not only shared across ploidy levels but was also the most widespread haplotype, found in 67 samples across 34 populations. This included geographically distinct species such as the Scottish endemic *E. marshallii* and the predominantly English and Welsh *E. anglica*, and ecologically contrasting taxa such as the dry heathland specialist *E. micrantha* and the (currently unpublished) obligate coastal “*E. fharaidensis*” Overall, 86% of taxa had one of six widespread haplotypes. There were also a large number of rare haplotypes, with over two thirds restricted to a single population (17 haplotypes: H4, H7-H10, H12-H21 and H23; Table 1). The remaining haplotypes found in multiple populations (H1, H2, H3, H5, H6, H11) showed no clear pattern of geography, with three found in all geographic regions (England, Scotland, Wales) and the remaining three shared between two geographic regions. Similarly, patterns of haplotype sharing do not follow species boundaries. Of the eight species with multiple populations (excluding hybrids), none of them had diagnostic ITS haplotypes. Despite haplotype sharing across taxa,there was no evidence for this being due to non-neutral processes, as the value of Tajima’s D (-0.17) was not significantly different from zero.

**Table 1.**
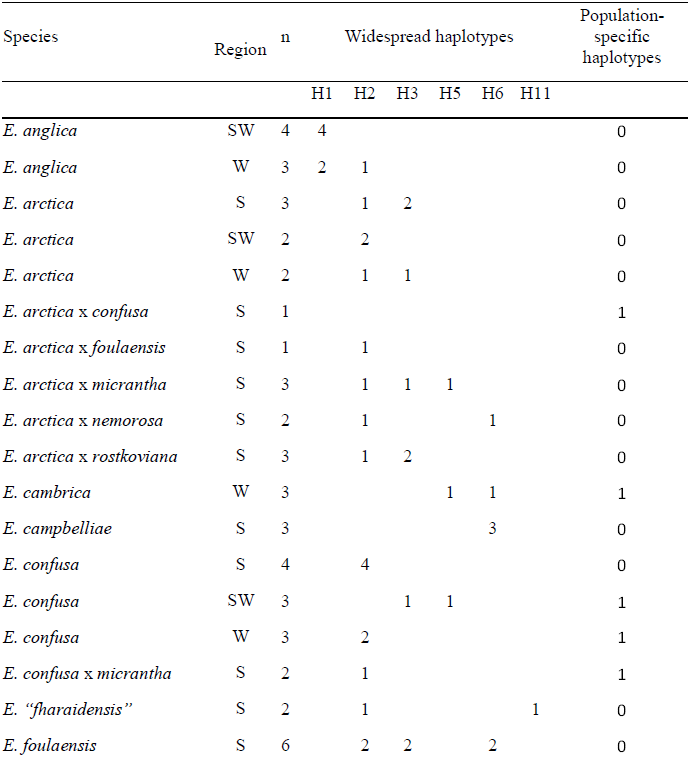
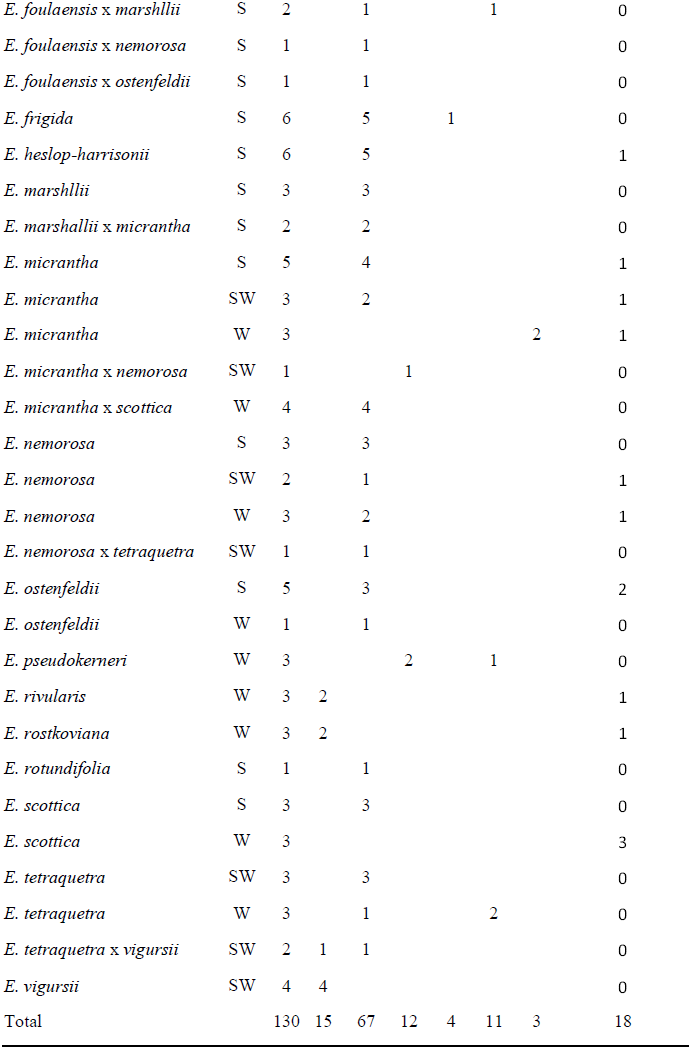
The distribution of ITS2 haplotypes between species and geographic regions in British *Euphrasia*. Haplotype numbers correspond to the haplotype network Figure 2. Population specific haplotypes are aggregated under one column. n = number of samples.

The putative hybrid species, *E. vigursii* and *E. rivularis*, possessed ITS haplotype H1, which is common to other diploid taxa, or population specific haplotypes (H20, H21), but no tetraploid haplotypes. The two sampled diploid-tetraploid hybrids (*E. arctica* x *rostkoviana*,*E. tetraquetra* x *vigursii*) possessed the full range of haplotypes: haplotype H2, which is common across ploidy levels, tetraploid specific haplotype H3, and diploid specific haplotype H1. Most (9/12) tetraploid hybrid populations had haplotypes shared with their putative parents, while the other populations had unique haplotypes.

The highest F_CT_ values in the SAMOVA were when *K* = 2 (Table S3 [**Supplementary Information**]), and this corresponded to the diploid-tetraploid divide described above. At *K*= 3, SAMOVA distinguished clusters corresponding to the two ploidy groups, and a third group of hybrid species derived from inter-ploidy level mating. AMOVA also supported the strong division by ploidy, with 88.2% of variation attributed to ploidy differences (Table 2; *P*< 0.001). A high proportion of variation was also partitioned by taxa (63.2%), and regions (25%), though this may be inflated by limited sampling within species.

**Table 2.** Hierarchical analysis of molecular variance (AMOVA) of British *Euphrasia* populations. Analyses performed between (A) species, (B) 3 geographic locations (Wales, South-West England, Scotland, (C) diploids and tetraploids. Number in parentheses are the results only including species (excluding hybrids). d.f. = Degrees of freedom. \*\**P* < 0.001;\**P* < 0.05.

| Source of variation | ITS |  | Plastid DNA |  |
| --- | --- | --- | --- | --- |
|  | d.f. | % Total variance | d.f. | % Total variance |
| **(A) Taxa** |  |  |  |  |
| Between taxa | 33 (19) | 63.17**<br>(65.62**) | 33 (19) | 15.87** (26.48**) |
| Within taxa | 96 (77) | 36.83 (34.38) | 96 (68) | 84.13 (73.52) |
| **(B) Location** |  |  |  |  |
| Between regions | 2 (2) | 25.52**<br>(25.92**) | 2 (2) | 5.40** (4.91*) |
| Within regions | 127 (94) | 74.48 (74.08) | 127 (85) | 94.60 (95.09) |
| **(C) Ploidy** |  |  |  |  |
| Between ploidy groups | 1 (1) | 88.24**<br>(84.39**) | 1 (1) | 11.05* (18.76*) |
| Within Diploid and tetraploid | 123 (95) | 11.76 (15.61) | 123 (86) | 88.95 (81.24) |

Summary statistics and haplotype networks revealed substantial divergence between diploid and tetraploid haplotypes. An average of 18.3 site differences were found between diploids and tetraploids, with divergence measured as D_XY_ = 0.051 (5.1%). The haplotype network revealed clusters corresponding to diploid and tetraploid haplotypes, separated by many mutations (Figure 2). The diploid cluster centres round haplotype H1, found in 15 individuals from 5 diploid species and one diploid-tetraploid hybrid. The only haplotype from this part of the network present in tetraploids is haplotype H18, found in a single sample of *E. ostenfeldii*. Within the tetraploid cluster, widespread haplotype H2 is at the centre, surrounded by other widespread haplotypes (H3, 8 populations, 12 samples; H6, 7 populations, 11 samples), and singleton haplotypes.

**Figure 2.**
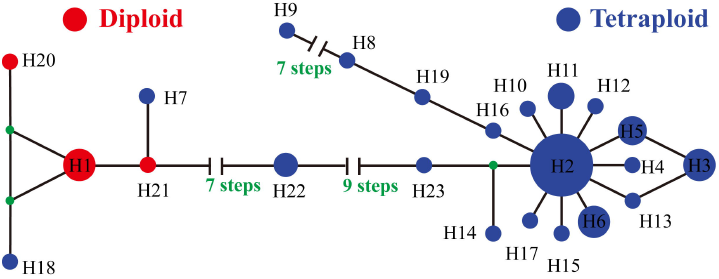
Median-joining network of ITS haplotype relationships in British *Euphrasia*. Numbers correspond to the ITS haplotypes in Table 1. Haplotypes are coloured by ploidy, with diploids in red and tetraploids in blue. Hypothetical (unsampled) haplotypes are represented by filled black circles.

### ITS phylogeny and molecular dating

Our broad-scale global *Euphrasia* phylogenetic analyses performed using MrBayes gave meaningful clusters of species, though the tree topology was generally poorly supported with many polytomies (Fig. 3). Haplotypes occurring in Britain were predominantly found in two main clusters: a tetraploid clade of Holarctic taxa from Sect. *Euphrasia* (posterior probability support, pp = 1.00, Clade A, Fig. 3), and a well-supported geographically restricted Palearctic diploid lineage (pp = 1.0, Clade B, Fig. 3). The tetraploid clade included a mix of British and European taxa, and is sister to a mixed clade of alpine diploid species and tetraploid *E. minima* (Clade IVc, Fig. 3). The diploid clade includes British diploids *E. anglica*, *E. rivularis*, *E. rostkoviana*, *E. vigursii* and European relatives (diploids or taxa without chromosome counts). The only non-diploid in the clade is one individual of tetraploid British *E. ostenfeldii*, which appears to be correctly identified and thus may have captured the diploid ITS haplotype through historical hybridisation. Overall, terminal branches of the tree are short, indicative of limited variation between related haplotypes. The only exception was the long branch of *E. disperma* from New Zealand, a result seen in previous Bayesian analyses (cf. Gussarova et al., 2008, fig 2) but not in Parsimony analyses, where it clusters together with the other southern hemisphere species on a shorter branch (Gussarova et al., 2008).Molecular dating with BEAST yielded a similar tree topology to MrBayes (results not shown), with many nodes having a low posterior support values. The clade of tetraploids and alpine relatives had an estimated crown age of 2.2 Ma (95% HPD = 2.0 - 2.4 Ma, Fig. 3, Node 1). The median crown age of the major group of diploids (also including Nearctic endemics *E. oakesii* and *E. randii*) was estimated as 1.0 Ma (95% HPD = 0.7 – 1.4 Ma; pp = 1.00, Fig. 3, Node 2). Due to low support of internal branches, the age of the most recent common ancestor of the diploid and tetraploid lineages could not be inferred directly. The nearest dated node gaining support is the broader clade of *Euphrasia*, which includes the diploid and tetraploid groups, in addition to two additional clades that include divergent species of unknown ploidy such as *E. insignis* (Japan), which has a crown age of 8.0 Ma, 95% HPD = 6.5 – 9.8 Ma (Fig. 3, Node 3).

**Figure 3.**
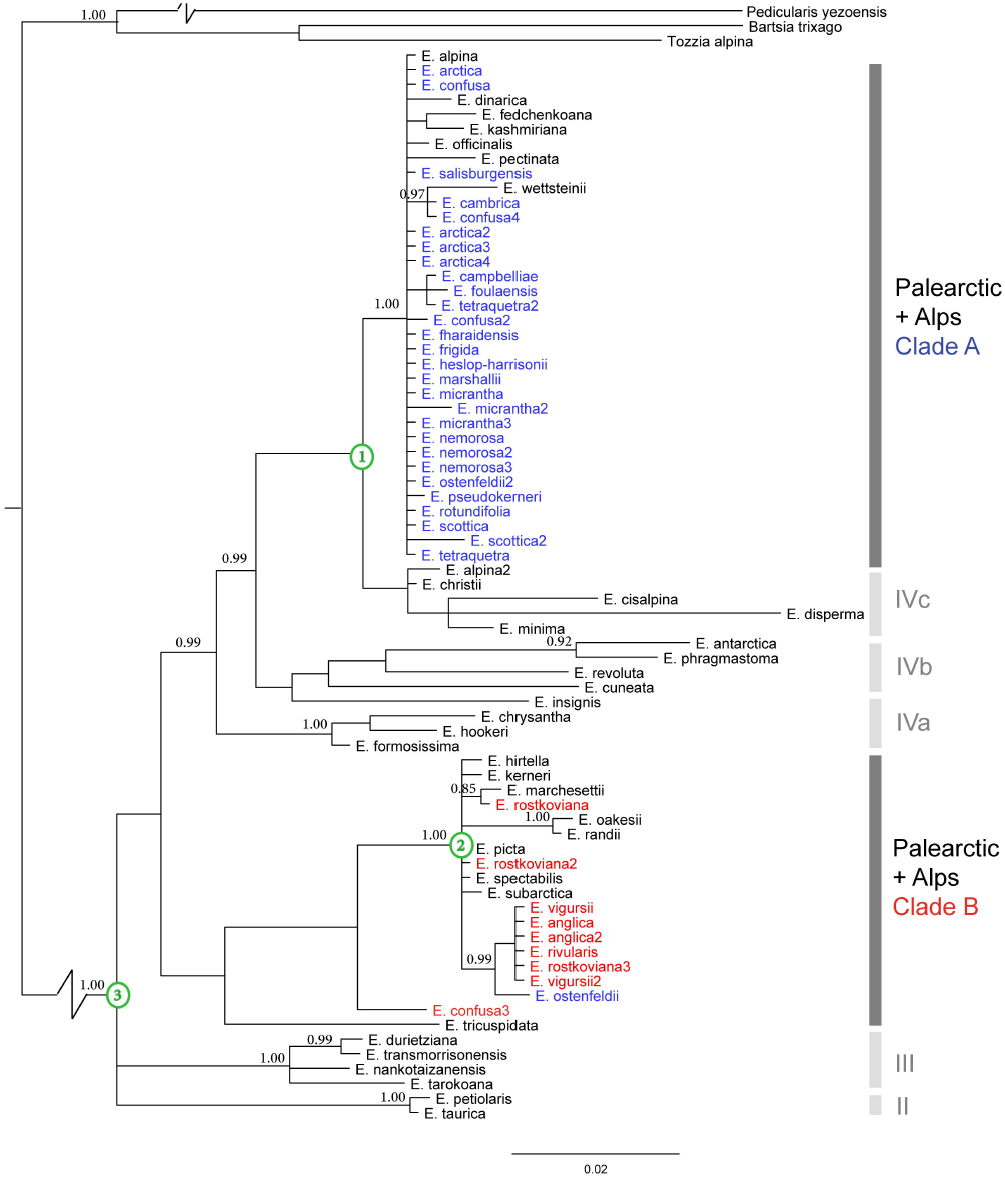
Majority rule consensus phylogeny of *Euphrasia* inferred from the ITS region using MrBayes. Posterior probabilities greater than 0.85 are indicated. Individuals are coloured by ploidy and geography: British diploids (red), British tetraploid (blue), other geographic areas (black). Green circled nodes have divergence dates estimated with BEAST, as given in the text. Clade A and Clade B correspond to the main study groups, with additional clades corresponding to Gussarova et al. (2008) also marked: II northern tetraploids; III Taiwan; IVa South American/Tasmanian; IVb complex (S. American, N. Zealand, Japan); IVc Alpine European.

### Plastid haplotype diversity

Initial sequencing of *rbcL* in 48 samples revealed no polymorphism, and so no further sequencing was performed for this region and it was excluded from further analyses.The final *matK* alignment was 844 bp with one indel, and the *rpl32-trnL* region was 630 bp with nine indels. The final concatenated plastid alignment was 1474 bp for 130 samples, with 2.7% segregating sites (40 sites) across 38 haplotypes. Nucleotide diversity was exceptionally low with Pi = 0.3%, and theta (per site) was similarly low at 0.00381. Tajima’s D was not significantly different from zero (-0.27).

Similar to ITS, most plastid haplotypes were individual or population specific (63%, 24/38 haplotypes found in one population only), with only 4 haplotypes being widespread (H4: 15 populations, 24 samples; H5: 13 populations, 18 samples; H2: 11 populations, 16 samples;H1, 10 populations, 14 samples, Table S4 [**Supplementary Information**]). However, unlike ITS, plastid haplotypes revealed complex patterns unrelated to ploidy. Most widespread haplotypes were shared across ploidy levels. An AMOVA found a moderate degree of genetic diversity was partitioned by ploidy (18.7%) and species (26.5%), with these values being reduced when hybrids were included (Table 2 [**Supplementary Information**]).Despite geography explaining little of the variation across the total dataset (4.9%), or being evident in the SAMOVA (Table S5), localised haplotype sharing was apparent in Scottish tetraploids. For example, haplotype H7 is shared across Scottish populations of *E. arctica* (and its hybrids), *E. foulaensis* and *E. micrantha* (and its hybrids), while H10 is also shared across three species in Scotland.

### Relationship among plastid haplotypes

The final concatenated plastid alignment was 1692 bp in length, for a total of 82 *Euphrasia* samples, including those from Gussarova et al. (2008). This alignment included the *trn*L intron (517 bp, 73 variable sites), *trn*L-*trn*F (420 bp, 85 variable sites) and *atp*B-*rbc*L (754 bp, 89 variable sites). The plastid tree (Fig. 4) successfully recovered the geographic clades reported in Gussarova et al. (2008). All diploid and tetraploid British samples possessed plastid haplotypes from the broad Palearctic taxa clade, which also includes *E. borneensis* (Borneo) and *E. fedtschenkoana* (Tian Shan). This clade received moderate support in our analysis (pp = 0.85). While informative of broad-scale relationships, most terminal branches were extremely short, and gave no information on interspecific relationships.

**Figure 4.**
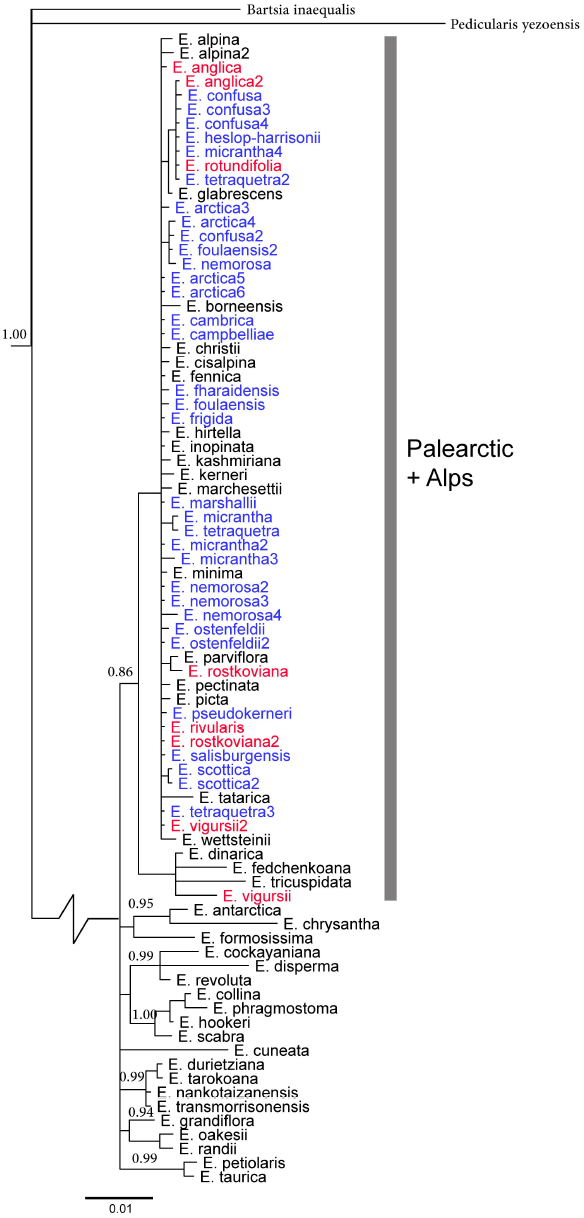
Majority rule consensus phylogeny of *Euphrasia* inferred from a concatenation of plastid *trn*L intron, *trn*L-*trn*F and *atp*B-*rbc*L using MrBayes. Posterior probabilities greater than 0.85 are indicated, and British diploids (red) and British tetraploid (blue) are coloured.

## DISCUSSION

We have investigated the utility of DNA barcoding data for the study of taxonomically complex *Euphrasia* in Britain. We find that *Euphrasia* species do not possess diagnostic ITS and plastid sequence profiles, and instead haplotypes are either widespread across taxa, or are individual or population specific. While DNA barcoding is of limited value as a molecular tool for identifying *Euphrasia* species, we show these data to be informative of the evolutionary processes underlying the generation and maintenance of diversity. Most notably, deep divergence of ITS haplotypes between ploidy groups (Pi = 5.1%, >2 Ma divergence), support British *Euphrasia* being assembled from diverse pre-glacial continental taxa, and with ploidy differences acting as an important reproductive barrier partitioning genetic variation. Overall our results shed light on the maintenance of genetic variation in one of the most renowned taxonomically challenging plant groups.

### DNA barcoding in taxonomically complex genera

Taxonomically complex *Euphrasia* have many characteristics that would make a DNA-based identification system desirable. In particular, molecular identification tools could be used to confirm species identities and subsequently revise the distribution of these under-recorded taxa. More generally, genetic data could be used to investigate which British species are genetically cohesive ‘good’ taxa. Our data show, however, that barcode sequences do not correlate with species boundaries as defined by morphology. Instead, haplotypes are often individual or population specific (for both ITS and plastid DNA), and where widespread are either shared across species within a ploidy level (for ITS) or across ploidy levels and species (plastid DNA). Patterns of haplotype sharing are often surprising, including between geographically disparate populations 100+ km apart, and between ecologically specialised taxa that seldom co-occur in the wild.

One possibility is that the species are not discrete genetic entities, and that the current taxonomy reflects a blend of discrete lineages, polytopic taxa, and morphotypes determined by a small number of genes. Alternatively, the species may represent meaningful biological entities, but with boundaries permeable to gene flow. In either case, the lack of species- specific or morphotype-specific barcodes may be affected by numerous evolutionary processes. Low sequence diversity associated with recent diversification is the first important factor. While elevated plastid diversity is a hallmark of some parasitic taxa, this is not the case for facultative hemiparasites like *Euphrasia*, which generally show similar patterns of mutation to autotrophic taxa (Wicke et al., 2016). Despite observing low plastid diversity (<0.5%) across taxa, we still detected 38 plastid haplotypes. This is sufficient variation for population genetic analysis, but lack of nucleotide diversity could explain the poor performance of our phylogenetic analyses. A second factor that reduces the ability of DNA data to distinguish species is selective sweeps, which can act on any genomic region including the plastid (Twyford, 2014, Muir and Filatov, 2007). While our data collection was not designed to measure the strength of selection, values for Tajima’s D were not significantly different from zero thus do not point to a strong selective regime.

In contrast to the factors above, it seems that self-fertilization, hybridity and incomplete lineage sorting are dominant factors shaping haplotype distributions across British *Euphrasia* species and populations. We observed many haplotypes that are restricted to individual samples or populations, consistent with French et al. (2008), who used extensive intrapopulation sampling to show most *Euphrasia* populations are fixed for a single plastid haplotype. This pattern of local fixation of haplotypes, and scarcity of widespread haplotypes, is likely due to many *Euphrasia* species being (at least partly) self-fertilising (Fis >0.4,French et al., 2004), and localised gravity-mediated seed dispersal. Where widespread haplotypes are present, these are shared amongst taxa and geographic areas. In these cases, dispersal of haplotypes has clearly occurred and their sharing among species may be due to hybridisation and incomplete lineage sorting. The large number of reported hybrids and hybrid species based on morphology (Preston and Pearman, 2015, Stace et al., 2015), as well as the prevalence of hybridisation in genetic data (Stone, 2013, Liebst, 2008), point to hybridisation being a key factor shaping genetic diversity in *Euphrasia*. Future genomic surveys will estimate the proportion of loci introgressing across species barrier in models that explicitly account for incomplete lineage sorting (Twyford and Ennos, 2012).

The sharing of DNA barcodes among *Euphrasia* species parallels a number of other studies where DNA barcoding has failed to provide species-level information. A notable example is willows, where hybridisation and selective sweeps have caused a single haplotype to spread across highly divergent taxa and between geographic regions (Percy et al., 2014). Poor species discrimination from DNA barcoding is also seen in the rapid radiation of Chinese *Primula* (Yan et al., 2015a) and *Rhododendron* (Yan et al., 2015b), which is likely a product of hybridisation and recent species divergence. In each of these groups, future DNA barcoding systems that target large quantities of nuclear sequence variation may provide resolution (Coissac et al., 2016, Hollingsworth et al., 2016). These data have the joint benefit of providing many nucleotide characters from unlinked loci, while also moving away from genomic regions that have atypical inheritance and patterns of evolution (i.e. plastids).Analyses of many nuclear genes (or entire genomes) would be particularly valuable for *Euphrasia*, where it may be possible to identify adaptive variants maintained in the face of hybridisation. These adaptive genes may underlie differences between species or ecotypes (Twyford and Friedman, 2015), and these loci could then potentially be used for future species identification.

### Polyploidy and the maintenance of genetic variation

The deep divergence of ITS haplotypes between diploid and tetraploid *Euphrasia* suggest strong reproductive barriers between ploidy levels, a result consistent with previous population surveys with AFLPs (French et al., 2008). The lack of support of internal nodes in our phylogeny make the divergence between ploidy groups difficult to date, but it must predate the origin of the tetraploid clade (2.2 Ma) and the diploid clade (1 Ma), with a date likely to be closer to 8 Ma (similar to a previous global *Euphrasia* phylogeny, Gussarova et al., 2008). Our divergence age estimate suggests that genetic diversity in British *Euphrasia* long pre-dates recent glacial divergence and the origin of young British endemic taxa. As such, British *Euphrasia* diversity has been assembled from a diverse pool of genetic diversity from European and Amphi-Atlantic taxa. The presence of distinct ITS haplotypes in each ploidy group may also be a consequence of concerted evolution of this multi-copy region (Sang et al., 1995), rather than absolute reproductive isolation. Further work will be required to test the extent of gene flow across ploidy levels, which is now emerging as a common feature of other plant groups (Pinheiro et al., 2010, Slotte et al., 2008).

The divergence between diploid and tetraploid ITS sequences adds further weight to the British tetraploids not being young autopolyploids, and instead being allopolyploids. While it is difficult to decipher the parentage of British tetraploids from our data, our phylogenetic analysis place these British taxa in a clade composed exclusively of tetraploids (Clade IVd, Gusarova et al., 2008), and thus these species may have originated elsewhere before dispersal to the UK. Alternatively, plastid haplotypes shared between diploids and tetraploids could point to a British diploid parent to the tetraploids, with this parent not contributing an ITS haplotype. An allotetraploid origin is further supported by the high number of tetraploid-specific AFLP bands (French et al., 2008), fixed microsatellite heterozygosity indicative of disomic inheritance (Stone, 2013), and tetraploid genome assemblies of double the size of diploids (Twyford and Ness, Unpublished data). Future genomic and cytogenetic studies will clarify the origin of British tetraploids.

## Conclusions

This study highlights how DNA barcoding data may fail to distinguish between species in taxonomically complex groups such as *Euphrasia*. No species in our study possessed a consistent diagnostic haplotype. Widespread haplotype sharing among species, in conjunction with high levels of intraspecific variation, make *Euphrasia* a particular challenge for DNA barcoding. However, our results are able to help us understand the maintenance of diversity, and in particular allow us to comment on the origins of British tetraploid species.

## ACKNOWLEDGEMENTS

Thanks to Richard Ennos for his assistance with this research. This study formed part of an international exchange programme for XW, funded by a Chinese Scholarship Council (CSC). The Royal Botanic Garden Edinburgh (RBGE) is supported by the Scottish Government’s Rural and Environmental Science and Analytical Services Division. Research by ADT is supported by NERC Fellowship NE/L011336/1.

---

**Supplementary Table S1.** Voucher information and population details of *Euphrasia* samples used in the study. Columns headed cpDNA and ITS 2 refer to the number of individuals sequenced for these regions.

**Supplementary Table S2.** PCR conditions and primer sequences for regions sequenced in this study.

**Supplementary Table S3.** Spatial analysis of molecular variation (SAMOVA) of ITS sequence data across British *Euphrasia* populations.

**Supplementary Table S4.** Plastid haplotype frequencies across British *Euphrasia* species. The column cpDNA indicates the number of individuals sampled.

**Supplementary Table S5.** Spatial analysis of molecular variation (SAMOVA) of plastid sequence data across British *Euphrasia* populations.

## LITERATURE CITED

Bandelt H-J, Forster P, Röhl A. 1999. Median-joining networks for inferring intraspecific phylogenies. Molecular Biology and Evolution, 16: 37–48.

Brysting AK, Oxelman B, Huber KT, Moulton V, Brochmann C, Renner S. 2007. Untangling complex histories of genome mergings in high polyploids. Systematic Biology, 56: 467–476.

Coissac E, Hollingsworth PM, Lavergne S, Taberlet P. 2016. From barcodes to genomes: extending the concept of DNA barcoding. Molecular Ecology, 25: 1423–1428.

de Vere N, Rich TC, Ford CR, Trinder SA, Long C, Moore CW, Satterthwaite D, Davies H, Allainguillaume J, Ronca S. 2012. DNA barcoding the native flowering plants and conifers of Wales. PLoS One, 7: e37945.

Dellicour S, Mardulyn P. 2014. SPADS 1.0: a toolbox to perform spatial analyses on DNA sequence data sets. Molecular Ecology Resources, 14: 647–651.

Dupanloup I, Schneider S, Excoffier L. 2002. A simulated annealing approach to define the genetic structure of populations. Molecular Ecology, 11: 2571–2581.

Drummond AJ, Suchard MA, Xie D & Rambaut A. 2012. Bayesian phylogenetics with BEAUti and the BEAST 1.7 Molecular Biology And Evolution 29: 1969–1973.

Ennos RA, French GC, Hollingsworth PM. 2005. Conserving taxonomic complexity. Trends in Ecology & Evolution, 20: 164–168.

Excoffier L, Lischer HE. 2010. Arlequin suite ver 3.5: a new series of programs to perform population genetics analyses under Linux and Windows. Molecular Ecology resources, 10: 564–567.

French G, Hollingsworth P, Silverside A, Ennos R. 2008. Genetics, taxonomy and the conservation of British *Euphrasia*. Conservation Genetics, 9: 1547–1562.

French GC, Ennos RA, Silverside AJ, Hollingsworth PM. 2004. The relationship between flower size, inbreeding coefficient and inferred selfing rate in British *Euphrasia* species. Heredity, 94: 44–51.

China Plant BOL Group. 2011. Comparative analysis of a large dataset indicates that internal transcribed spacer (ITS) should be incorporated into the core barcode for seed plants. Proceedings of the National Academy of Sciences, 108: 19641–19646.

Gussarova G, Popp M, Vitek E, Brochmann C. 2008. Molecular phylogeny and biogeography of the bipolar *Euphrasia* (Orobanchaceae): Recent radiations in an old genus. Molecular Phylogenetics and Evolution, 48: 444–460.

Hebert PD, Gregory TR. 2005. The promise of DNA barcoding for taxonomy. Systematic Biology, 54: 852–859.

Hodges SA, Arnold ML. 1994. Columbines: a geographically widespread species flock. Proceedings of the National Academy of Sciences, 91: 5129–5132.

Hollingsworth P, De-Zhu L, Van der Bank M, Twyford A. 2016. Telling plant species apart with DNA: from barcodes to genomes. Philosophical Transactions of the Royal Society B.

Hollingsworth PM, Forrest LL, Spouge JL, Hajibabaei M, Ratnasingham S, van der Bank M, Chase MW, Cowan RS, Erickson DL. et al. 2009. A DNA barcode for land plants. Proceedings of the National Academy of Sciences, 106: 12794–12797.

Hollingsworth PM, Graham SW, Little DP. 2011. Choosing and using a plant DNA barcode. PLoS ONE, 6: e19254.

Kolář F, Píšová S, Záveská E, Fér T, Weiser M, Ehrendorfer F, Suda J. 2015. The origin of unique diversity in deglaciated areas: traces of Pleistocene processes in north-European endemics from the *Galium pusillum* polyploid complex (Rubiaceae). Molecular Ecology, 24: 1311–1334.

Librado P, Rozas J. 2009. DnaSP v5: a software for comprehensive analysis of DNA polymorphism data. Bioinformatics, 25: 1451–1452.

Liebst B. 2008. Do they really hybridize? A field study in artificially established mixed populations of *Euphrasia minima* and *E. salisburgensis* (Orobanchaceae) in the Swiss Alps. Plant Systematics and Evolution, 273: 179–189.

McNeal JR, Bennett JR, Wolfe AD, Mathews S. 2013. Phylogeny and origins of holoparasitism in Orobanchaceae. American Journal of Botany, 100: 971–983.

Muir G, Filatov D. 2007. A selective sweep in the chloroplast DNA of dioecious *Silene* (Section Elisanthe). Genetics, 177: 1239–1247.

Percy DM, Argus George W, Cronk Quentin C, Fazekas Aron J, Kesanakurti Prasad R, Burgess Kevin S, Husband Brian C, Newmaster Steven G, Barrett Spencer CH, Graham Sean W. 2014. Understanding the spectacular failure of DNA barcoding in willows (*Salix*): Does this result from a trans-specific selective sweep? Molecular Ecology, 23: 4737–4756.

Pinheiro F, De Barros F, Palma-Silva C, Meyer D, Fay MF, Suzuki RM, Lexer C, Cozzolino S. 2010. Hybridization and introgression across different ploidy levels in the Neotropical orchids *Epidendrum fulgens* and *E. puniceoluteum* (Orchidaceae). Molecular Ecology, 19: 3981–3994.

Preston CD, Pearman DA. 2015. Plant hybrids in the wild: evidence from biological recording. Biological Journal of the Linnean Society, 115: 555–572.

Sang T, Crawford DJ, Stuessy TF. 1995. Documentation of reticulate evolution in peonies (*Paeonia*) using internal transcribed spacer sequences of nuclear ribosomal DNA:implications for biogeography and concerted evolution. Proceedings of the National Academy of Sciences, 92: 6813–6817.

Schmickl R, Paule J, Klein J, Marhold K, Koch MA. 2012. The evolutionary history of the *Arabidopsis arenosa* complex: diverse tetraploids mask the western carpathian center of species and genetic diversity. PLOS ONE, 7: e42691.

Slotte T, Huang H, Lascoux M, Ceplitis A. 2008. Polyploid speciation did not confer instant reproductive isolation in *Capsella* (Brassicaceae). Molecular Biology and Evolution, 25: 1472–1481.

Spooner DM. 2009. DNA barcoding will frequently fail in complicated groups: An example in wild potatoes. American Journal of Botany, 96: 1177–1189.

Squirrell J, Hollingsworth PM, Bateman RM, Tebbitt MC, Hollingsworth ML. 2002. Taxonomic complexity and breeding system transitions: conservation genetics of the *Epipactis leptochila* complex (Orchidaceae). Molecular Ecology, 11: 1957–1964.

Stace CA, Preston CD, Pearman DA. 2015. Hybrid flora of the British Isles: Botanical Society of Britain and Ireland.

Stone H. 2013. Evolution and conservation of tetraploid Euphrasia L. in Britain. Unpublished Thesis, The University of Edinburgh.

Svensson BM, Carlsson BÅ. 2004. Significance of time of attachment, host type, and neighbouring hemiparasites in determining fitness in two endangered grassland hemiparasites. Annales Botanici Fennici, 41: 63–75.

Young ND, Healy J. 2003. GapCoder automates the use of indel characters in phylogenetic analysis. BMC Bioinformatics, 4: 6.

